# X-chromosomal diversity may, or may not, reflect climate

**DOI:** 10.1101/2023.09.15.557915

**Authors:** Zarus Cenac

**Affiliations:** Independent Researcher

## Abstract

Previous research indicates that a climatic signal is present in mitochondrial diversity, but absent in other genetic diversities (e.g., X-chromosomal) and cranial diversity. Such research included analysis which adjusted diversity for distance from Africa (i.e., to account for global expansion), and saw whether adjusted diversity is associated with climate. The indication of a climatic signal may be affected by the choice of African location from which distance is measured. To bypass this potential effect of location, some analyses in the present research only featured populations which are located outside of Africa. The present research (Studies 1 and 2) used various diversities which were sourced from previous research. Autosomal, X-chromosomal, mitochondrial, and cranial diversities were adjusted for distance from Africa. This adjustment was not used regarding Y-chromosomal diversity; a preprint suggested that Y-chromosomal diversity does not signify expansion from Africa. In Study 1, adjusted X-chromosomal diversity increased with minimum temperature. Conscious of replicability, Study 2 examined whether a climatic signal is present in X-chromosomal diversity when an alternative dataset is used; a signal was not detected. Other adjusted diversities, including (surprisingly) mitochondrial, were not found to be associated with minimum temperature. Data from one dataset, but not another, was supportive of a relationship between sex-biased migration and climate—it seems unclear if this relationship would underlie a climatic signal in X-chromosomal diversity. Therefore, it is uncertain i) if X-chromosomal diversity is associated with climate, and, ii) why there might be an association. Further replication research on this topic is called for.

## Introduction

### Climate and global expansion

Research has considered factors which may affect human biological diversity, e.g., by examining expansion from Africa and climate (e.g., Balloux et al., 2009; Betti et al., 2012). In modern humans, population-level diversity falls as geographical distance from Africa rises (Betti et al., 2012; Prugnolle et al., 2005; von Cramon-Taubadel & Lycett, 2008) which indicates a worldwide expansion (through population bottlenecks) from Africa (Manica et al., 2007; Ramachandran et al., 2005; von Cramon-Taubadel & Lycett, 2008; see Figure 1). As for climate, mitochondrial DNA is known to have been affected by climate (Balloux et al., 2009); mitochondria are acknowledged to play a role in climatic adaptation (Grover-Thomas et al., 2026). Indeed, in Balloux et al. (2009), climate (minimum temperature) was associated with mitochondrial diversity. This association persisted when mitochondrial diversity was adjusted for linear distance from Africa—adjusted mitochondrial diversity rose with minimum temperature (Balloux et al., 2009; see Supplementary Note 1). No such association was apparent for adjusted autosomal, X-chromosomal, or Y-chromosomal diversities (Balloux et al., 2009) (see Supplementary Note 2). The shape diversity of the femur and tibia are related to minimum temperature (Betti et al., 2012), whilst a relationship with climate fails to be supported when it comes to cranial form diversity (Betti et al., 2009; see Supplementary Note 3) and pelvic shape diversity (Betti et al., 2012). Therefore, a climatic signal seems to be evident in some biological diversities (Balloux et al., 2009; Betti et al., 2012) yet not be evident in others (Balloux et al., 2009; Betti et al., 2009, 2012).

**Figure 1.**
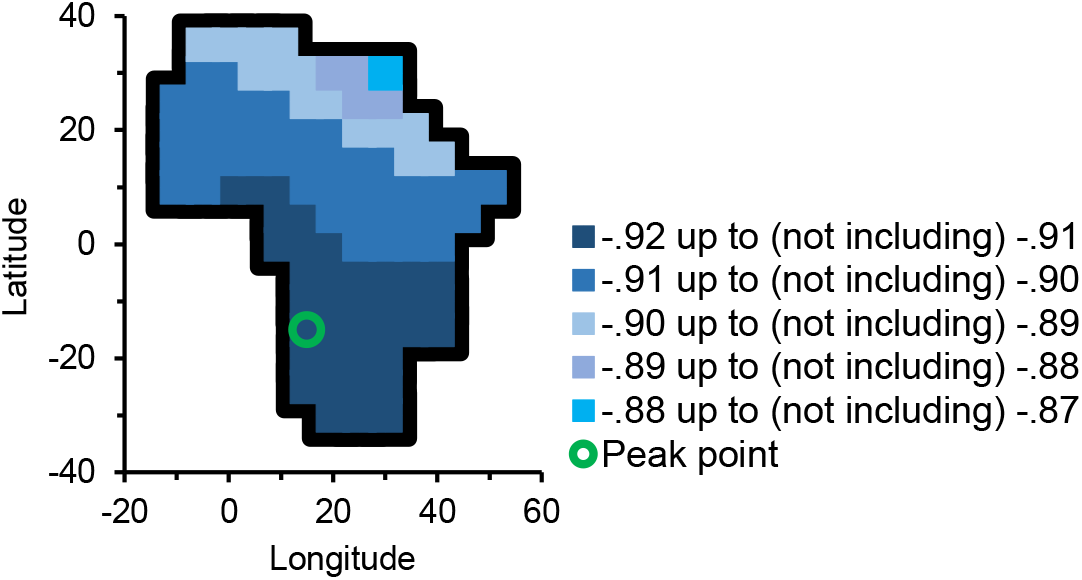
X-Chromosomal Diversity and the African Origin of Global Expansion. *Note*. In order to indicate where expansion started, various locations can be used (one at a time) as if each was the origin, and it can be seen how well distance from that location is related to diversity (e.g., Manica et al., 2007). Support for a location being the origin of expansion is indicated by the (relative) extent to which diversity declines as distance from a location increases—a stronger decline (and therefore a stronger expansion signal) bodes well for a location being the origin (e.g., Ramachandran et al., 2005; von Cramon-Taubadel & Lycett, 2008). The location which results in the most negative decline can be called the *peak point* (e.g., Cenac, 2022). Like has been done with correlation coefficients for relationships between genetic diversity and distance from Africa (Luca et al., 2011; Ramachandran et al., 2005), Figure 1 presents correlation coefficients (*r*—blue areas) regarding X-chromosomal microsatellite heterozygosity and distance from Africa (coefficients are from Cenac, 2023a, who used diversities from Balloux et al., 2009, and distances in Cenac, 2022). The *peak point* is the location giving the lowermost correlation coefficient (Cenac, 2022) (green circle). Figure 1 adjusts Figure 4 in Cenac (2023b) which employed coordinates (in Africa) that are contained in a figure by Betti et al. (2013) (Figure 4 in Cenac, 2023b, did not feature correlation coefficients, and it had some basis in Cenac, 2022).

### Diversity adjusted for distance

When studying whether diversity is related to climate, diversity can be adjusted for distance from Africa, and it can be seen if this adjusted diversity is related to climate (Balloux et al., 2009). However, when using populations pulled from across the world, the relationship between diversity (adjusted for linear distance from Africa) and climate can be affected by the location in Africa which distances are measured from (Figure 2A and 2C). Therefore, a supposed lack of relationship with climate (e.g., for adjusted X-chromosomal diversity in Balloux et al., 2009) could be a product of analysis having used distances from a location which just so happens to lead to a weaker correlation coefficient.

**Figure 2.**
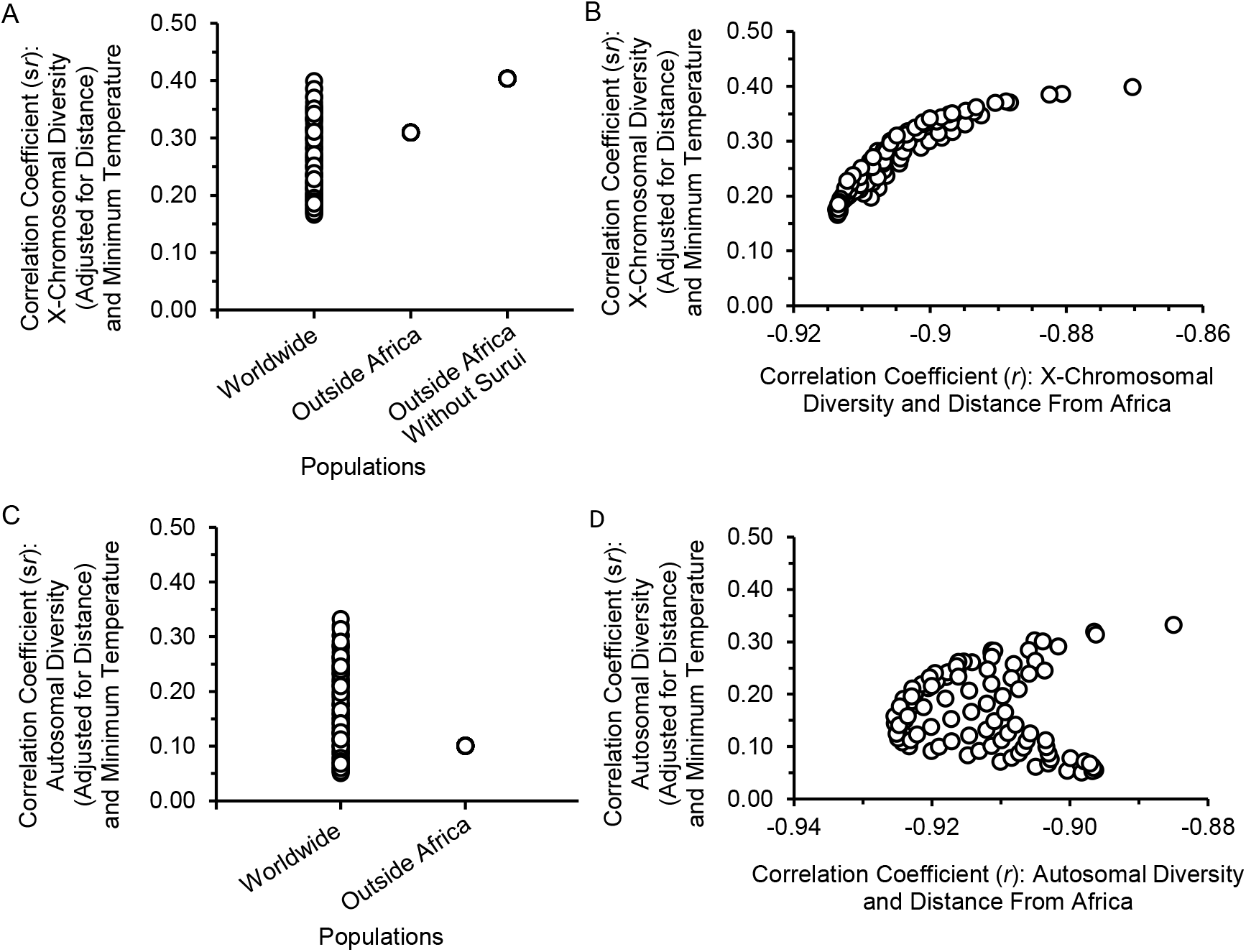
Diversity, Climate, and Distance from Africa. *Note*. Figure 2 used distances from 99 locations in Africa (Cenac, 2022), X-chromosomal microsatellite heterozygosities, autosomal single nucleotide polymorphism (SNP) haplotype heterozygosities, and minimum temperatures (Balloux et al., 2009). Figure 2A and 2B are with respect to X-chromosomal diversity, whilst Figure 2C and 2D are regarding autosomal diversity. In some instances, like in Balloux et al. (2009), diversity was adjusted for linear distance from Africa (see the *y*-axes of 2A, 2B, 2C, and 2D). With respect to minimum temperature (Balloux et al., 2009) and adjusted diversity, semi-partial Pearson correlation coefficients were calculated (*y*-axes). This procedure was done with and without populations in Africa. For X-chromosomal diversity, in the absence of the populations in Africa (Figure 2A), there was an atypical datapoint. The Surui population in the HGDP-CEPH data (Balloux et al., 2009) was this datapoint. Regarding populations from across the world, Balloux et al. used distance from a particular location in Africa—the present research observed quite similar *R*^2^s (compared to Balloux et al.) when using distances in the vicinity of that particular location— the corresponding s*r*s are generally amongst the lowest when populations are from across the world in Figure 2A and 2C. For Figure 2B and 2D, populations from across the world were used—a significance test was not run (there likely would be nonindependence—see Judd et al., 2009, regarding nonindependence), although patterns do seem to be visually indicated. For some diversities, a weaker correlation coefficient may indicate that the signal of expansion from Africa has been controlled for better—when it comes to X-chromosomal diversity, it appears that correlation coefficients may become weaker as more of the expansion signal is removed (Figure 2B). However, this is likely not the case for all diversities as it seems to not be the case for autosomal SNP haplotype heterozygosity (Figure 2D).

Distances from *which* location should be used to adjust diversity? The location whose distances deliver the strongest fall of diversity (e.g., Betti et al., 2013; Ramachandran et al., 2005) has been called the *peak point* (Cenac, 2022). The magnitude of the decline in diversity indicates the strength of the expansion signal (Ramachandran et al., 2005; von Cramon-Taubadel & Lycett, 2008). Therefore, to remove the influence of expansion on diversity, it would make sense to adjust diversity for distance from the peak point. However, different diversities can have different peak points (Cenac, 2022); did Balloux et al. (2009) use distance from diversity-specific peak points when they adjusted diversities? Balloux et al. (2009) used distance from a location (in Africa) from Manica et al. (2007). That location was arrived at through autosomal microsatellite heterozygosity and cranial form diversity (Balloux et al., 2009; Manica et al., 2007). The location would seem to be reasonably similar to the peak points for mitochondrial diversity and X-chromosomal diversity (in Cenac, 2022), less similar for the peak point (linear trend) for autosomal microsatellite heterozygosity (in Ramachandran et al., 2005), even less similar for autosomal SNP haplotype heterozygosity, but markedly different for Y-chromosomal microsatellite heterozygosity (in Cenac, 2022). Indeed, whilst peak points for autosomal, mitochondrial, and X-chromosomal diversity are in Africa (Cenac, 2022; Ramachandran et al., 2005), the peak point for Y-chromosomal microsatellite heterozygosity is not (Cenac, 2022).

Indeed, whereas Y-chromosomal microsatellite heterozygosity does decrease linearly as distance from Africa rises (Balloux et al., 2009), the strongest decline is not found when distance is from Africa; the peak point is located in Asia—Y-chromosomal microsatellite heterozygosity does not appear to portray expansion worldwide from Africa (Cenac, 2022). Therefore, when exploring if Y-chromosomal microsatellite heterozygosity is associated with climate, rather than adjusting diversity for distance from Africa (Balloux et al., 2009), it *may* make more sense to adjust diversity for distance from Asia. Nevertheless, global expansion from Asia seems unlikely (e.g., Li et al., 2008; Ramachandran et al., 2005). Therefore, all in all, Balloux et al. (2009) are likely to have greatly adjusted diversities for a global expansion signal except when it came to Y-chromosomal diversity.

Nonetheless, regarding populations who are located across the world, peak points (when they are in Africa) appear to be greatly determined by African populations (Cenac, 2022). This is problematic because few African populations *may* feature in data which are used in the calculation of a peak point (e.g., seven African populations regarding X-chromosomal and autosomal haplotype diversity) (Cenac, 2022). And so, it seems unclear if it is wise to simply use the peak point when diversity is adjusted for distance.

Therefore, when adjusting diversity, there is ambiguity concerning which location the distances should be from. An alternative approach could be to only use populations outside of Africa—when only those populations are used, the correlation coefficient does not vary according to the location in Africa that is used as the origin (Figure 2A and 2C). However, to comprehensively understand if there is a climatic signal in diversity, it would not be desirable to exclude an entire continent.

### Cranial form diversity

Betti et al. (2009) explored climate and diversity in a different way than Balloux et al. (2009) did. Betti et al. used coordinates across the Earth like each was an origin. For each origin, they started with a model for predicting cranial form diversity, using distance (to populations) terms and climatic terms. For each origin, Betti et al. (2009) found the most effective model for predicting diversity. Out of all the origins, the origin which had a model that best predicted diversity was an origin in Africa, and climate was not included in that model (Betti et al., 2009). This finding happened with respect to male crania, and female crania too (Betti et al., 2009). However, as mentioned above, the peak point may be particularly influenced by African populations (Cenac, 2022), which could be worth considering in the event that models using other *origins* included climatic terms in Betti et al. (2009).

### The present research

In two studies, it was examined whether diversities appear to have a climatic signal when the signal of expansion is controlled for (see Supplementary Note 4). Prior research has used minimum temperature as a climatic variable (Balloux et al., 2009; Betti et al., 2009, 2012; Manica et al., 2007; see Supplementary Note 5); minimum temperature was used as such in the present research. Using populations from across the world, earlier research adjusted genetic diversities for distance from Africa (distance from the same location in Africa for each diversity), and examined if these adjusted diversities are related to minimum temperature (Balloux et al., 2009); this approach was broadly taken in the present research, but with some differences. To counter a potential influence of African location on whether relationships with climate are apparent (e.g., Figure 2), several analyses solely featured populations outside of Africa. Those analyses concerned autosomal microsatellite heterozygosity, autosomal SNP haplotype heterozygosity, X-chromosomal microsatellite heterozygosity, and cranial form diversity. Certain analyses (global/worldwide analyses) featured African populations. Distances from the peak point specific to the relevant diversity (Cenac, 2022, 2023a) were used in the global analyses. These analyses featured the aforementioned autosomal and X-chromosomal diversities, as well as (for completeness) Y-chromosomal microsatellite heterozygosity.

A climatic signal in mitochondrial diversity has previously been indicated in *two* datasets (Balloux et al., 2009); exploring replicability in science is important (e.g., Nosek & Errington, 2020) and, in the present research, a second study (Study 2) was undertaken as a follow-up to Study 1 in order to further test whether a climatic signal in X-chromosomal diversity is indicated by using an alternative dataset (see Balloux et al., 2009; Mallick et al., 2016).

In the prior study which found that mitochondrial diversity (adjusted for distance from Africa) is associated with minimum temperature, the two datasets with which that observations were found were the mtDB and HVRBase++ (Balloux et al., 2009); the present research (in Study 2) sought to find whether those findings are replicated when mitochondrial diversity from the HGDP-CEPH (Lippold et al., 2014) is used.

Research has occurred with respect to sex-biased migration in human history (e.g., Keinan & Reich, 2010; Rasteiro et al., 2012). As a way of indicating whether a relationship between sex-biased migration and climate may underlie the climatic signal in mitochondrial diversity, previous research examined whether the ratio in diversity of the sex chromosomes is related to minimum temperature (Balloux et al., 2009); using data from the HGDP-CEPH (Balloux et al., 2009), Study 2 examined whether the ratio (after adjusting X-chromosomal diversity for distance from Africa) is related to minimum temperature, and this was done for the purpose of exploring whether sex-biased migration and climate are related because their relationship could provide a reason for why, in Study 1, X-chromosomal diversity was related to climate. Sex-biased migration could be reflected in the ratio of X-chromosomal diversity to autosomal diversity (Hammer et al., 2008). Therefore, using data from the Simons Genome Diversity Project (Mallick et al., 2006), as a post-hoc follow-up to analysis with the ratio of X- to Y-chromosomal diversity, it was seen whether the ratio of X-chromosomal to autosomal diversity is related to climate.

## Method

### Study 1: Genetic and cranial diversities

#### Data

Study 1 used genetic diversities in Balloux et al. (2009). These diversities were heterozygosities (51 populations from the HGDP-CEPH) regarding autosomal microsatellites, autosomal SNP haplotypes, X-chromosomal microsatellites, and Y-chromosomal microsatellites (Balloux et al., 2009). Balloux et al. (2009) acquired the haplotype diversities from Li et al. (2008). Additionally, the form diversity of male crania (90 populations) and female crania (34 populations) outside of Africa (Betti et al., 2009) was used (regarding cranial data, Betti et al., 2009, cite Manica et al., 2007).

Geographical distances were from prior research (Betti et al., 2009; Cenac, 2022), some of which (in Cenac, 2022) included distances from i) 99 locations in Africa from Betti et al. (2013), and ii) 32 global locations found in von Cramon-Taubadel and Lycett (2008). Geographical distances (Betti et al., 2009; Cenac, 2022) from peak points (Betti et al., 2009; Cenac, 2022, 2023a) were used to adjust diversities. These distances (Betti et al., 2009; Cenac, 2022) were from peak points which are i) in Africa for cranial form diversity (Betti et al., 2009), X-chromosomal microsatellite heterozygosity, autosomal SNP haplotype heterozygosity (Cenac, 2022), and autosomal microsatellite heterozygosity (Cenac, 2023a), and (for completeness) ii) in Asia for Y-chromosomal microsatellite heterozygosity (Cenac, 2022). Because peak points can vary between diversities (Cenac, 2022), the distances (from peak points) used for adjusting diversities were the ones relevant/specific to the type of diversity. Distances from peak points were utilised even when analyses only consisted of populations outside Africa (despite reasoning in the *Introduction*). For peak point coordinates, see Table S1 (or references given in Table S1). Distances regarding HGDP-CEPH populations are in Table S3.

Regarding genetic data, it was known from beforehand which continent populations are located within (Cenac, 2022); it was clear which populations are inside Africa, and which are not. Labelling employed in Betti et al. (2009) was used to identify which populations featured in their study are outside of Africa.

Minimum temperatures shown in previous research (Balloux et al., 2009; Betti et al., 2009) were used (both Balloux et al., 2009, and Betti et al., 2009, sourced minimum temperatures from WORLDCLIM, and they cited Hijmans et al., 2005, with respect to WORLDCLIM).

#### Analysis

Analysis took place in R Version 4.0.5 (R Core Team, 2021) and Microsoft Excel. ppcor (Kim, 2015) was used for running semi-partial correlation tests, except for the semi-partial Pearson correlation tests used regarding autosomal microsatellite heterozygosity, and the semi-partial Spearman correlation test employed regarding the diversity of male crania.

Autosomal microsatellite heterozygosity may *appear* to fall linearly when distance from Africa increases (Prugnolle et al., 2005), but this type of diversity aligns more strongly with a non-linear trend (quadratic) than a linear decline (Cenac, 2023a). Therefore, autosomal microsatellite heterozygosity was adjusted for a quadratic (rather than a linear) relationship with distance from Africa. It was then seen if the adjusted autosomal diversity is related to minimum temperature. Semi-partial Pearson correlation coefficients concerning adjusted autosomal diversity and minimum temperature were calculated (for populations globally, and only outside Africa). The correlation coefficients were converted to *t*-statistic values using a formula in Kim (2015). A *p*-value can be found for a *t*-statistic value (e.g., Kim, 2015), and *p*-values were indeed calculated for the converted *t*-values.

In Betti et al. (2009), there was a non-linear association between distance from Africa and cranial form diversity for male crania (105 populations from across the globe). The model used in Betti et al. (2009) for predicting diversity from distance from the peak point had an intercept, a linear term, and a cubic term (see Supplementary Note 6); a variant of their model was used to calculate residuals in the present study (the residuals being diversity adjusted for distance from Africa)—the same distances were used as in Betti et al., an intercept was also utilised, as was a linear term, and also a cubic term when trying to predict diversity from distance, but populations were solely outside of Africa. The semi-partial Spearman correlation coefficient for adjusted cranial diversity and minimum temperature was determined. The formula in Kim (2015) was used for converting the correlation coefficient to a *t*-value, for which (like with autosomal diversity) a *p*-value was calculated.

Linear declines (with increasing distance from Africa) are shown in autosomal SNP haplotype heterozygosity and X-chromosomal diversity (Balloux et al., 2009), with a quadratic relationship attaining no support (over a linear relationship) (Cenac, 2023a). And so, adjusting those diversities for linear distance from Africa (e.g., Balloux et al., 2009) would seem to be acceptable. Therefore, analysis in the present study set out to adjust those diversities for linear distances.

Y-chromosomal microsatellite heterozygosity has a peak point in Asia (and indicates an origin of expansion which possibly is exclusive to Asia), with a linear decline from there (Cenac, 2022), and an absence of support for a non-linear relationship (Cenac, 2023a). Therefore, it was planned, for the present study, to adjust Y-chromosomal diversity for linear distance from Asia (whereas Balloux et al., 2009, adjusted for linear distance from Africa).

There are various ways of assessing whether datapoints are atypical (e.g., Field, 2013; Judd et al., 2009). In the present study, datapoints which have *z*-scores of residuals over |3.29| were (conservatively) noted as being atypical (e.g., Field, 2013). The Surui population (Balloux et al., 2009) had an atypical standardised residual in some analyses (see *Results*). In the absence of atypical datapoints, it was seen whether heteroscedasticity was indicated. It was also seen if the presence of heteroscedasticity was resolved in the absence of any one population. To visually assess whether heteroscedasticity was present (in parametric analysis), adjusted diversity (*y*-axis) was placed against minimum temperature (*x*-axis) in graphs. Several graphs of this sort are presented in Balloux et al. (2009).

To see whether positive spatial autocorrelation was present amongst residuals, the three-step method of Chen (2016) was employed. This was done utilising a spreadsheet which is available in Chen (2016). Spatial Durbin-Watson values (spatial *DW*s) are produced in the three-step method, and Durbin-Watson bounds are applicable to the spatial *DW* (Chen, 2016). Consequently, 5% Durbin-Watson bounds (Savin & White, 1977) were referred to in order to assess if positive spatial autocorrelation was at hand. However, spatial autocorrelation would enhance the Type I error rate (e.g., Deblauwe et al., 2012), therefore, when significant relationships were not evident (e.g., Figure 3), positive spatial autocorrelation would not *need* to be tested for.

**Figure 3.**
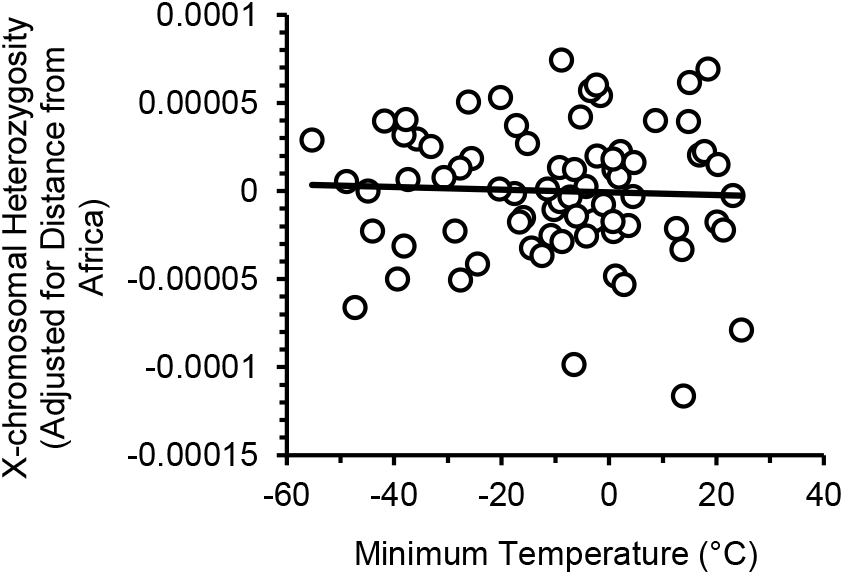
A Correlation was not Observed Between Adjusted X-Chromosomal Diversity and Minimum Temperature in Study 2. *Note*. The values of the datapoints are given in Table S2 as are the geographical distances (from Africa) which were used to adjust X-chromosomal diversity.

The Bonferroni correction (reviewed in Abdi, 2010, alongside other methods of adjusting *p*-values) was used on *p*-values in the current study; *p*-values were multiplied by five (as there were five types of diversity of interest in this study) regarding the genetic diversities, but multiplied by 10 concerning cranial diversity (multiplied by five given the five types of diversity, and multiplied by two because there were separate analyses of males and females).

### Study 2: Diversity and diversity ratios

#### Data

Study 2 explored whether there is a climatic signal in X-chromosomal diversity. The study used X-chromosomal expected heterozygosity (calculated from base pair dissimilarities) of persons whose sex is female (77 populations outside of Africa) from the Simons Genome Diversity Project (SGDP) (Mallick et al., 2016). The procedure of Williams (2011) was used for the purpose of calculating population geographical distances (from Africa). To calculate those distances, population coordinates (Mallick et al., 2016), waypoints (Cenac, 2022; von Cramon-Taubadel & Lycett, 2008), and an origin location in Africa (von Cramon-Taubadel & Lycett, 2008) were used. Minimum temperatures (each month from 1981 to 2025) were obtained from NASA POWER (2026) using the population coordinates. The downloaded data (NASA POWER, 2026) included a column featuring the lowest minimum temperature (from the 12 months) for each year. These lowest minimum temperatures were averaged (across the years 1981 to 2025) in order to arrive at the *minimum temperature variable* which was used in the current study.

In Study 2, it was examined if mitochondrial diversity (adjusted for distance from Africa) is correlated with minimum temperature. Distance was from the same origin location (von Cramon-Taubadel & Lycett, 2008) used regarding X-chromosomal diversity in Study 2, and that location was selected for mitochondrial diversity for the same reason it was selected for X-chromosomal diversity (see the note in Table S3). Mitochondrial nucleotide diversity (44 populations) from HGDP-CEPH data (Lippold et al., 2014) was used, as was the minimum temperature for each population (Balloux et al., 2009). Those 44 populations are ones who are located outside of Africa (Cenac, 2022). When using mitochondrial diversity, which was calculated “as average pairwise difference” (Balloux et al., 2009, p. 3448), the change in mitochondrial diversity with distance from Africa has a quadratic trend (Cenac, 2023a). The mitochondrial data with which that trend was observed was data which Balloux et al. (2009) found in the mtDB, and the trend was found when analysis used populations gathered from inside and outside of Africa (Cenac, 2023a). Regarding the mtDB data, whether there was a quadratic trend was tested using the Bayesian information criterion (BIC) to compare between models (linear vs. quadratic) (Cenac, 2023a); in Study 2, when that methodology was applied to the HGDP-CEPH data, BICs did not suggest support for a quadratic model over a linear one (the difference in BICs was only .72).

Additionally, using HGDP-CEPH data (X- and Y-chromosomal diversities) and minimum temperatures (Balloux et al., 2009), Study 2 expanded upon Balloux et al. (2009) who looked into whether the ratio between the diversities of sex chromosomes is related to minimum temperature; the present study adjusted X-chromosomal diversity for distance from Africa before dividing the result by Y-chromosomal diversity—it was then seen if the resulting ratio correlated with minimum temperature. Regarding HGDP-CEPH data, geographical distances from a preprint (Cenac, 2022) were used.

Moreover, to explore whether a relationship between sex-biased migration and climate is indicated in another dataset (see Balloux et al., 2009; Mallick et al., 2016), X-chromosomal and autosomal heterozygosities from the SGDP (75 populations) (Mallick et al., 2016) were used to calculate a ratio from the two diversities (X divided by autosomal), and it was evaluated whether that ratio correlates with minimum temperature. Diversities were analysed without being adjusted for distance from Africa to avoid complications from using positive and negative numbers on the numerator and denominator of the ratio.

#### Analysis

Analysis utilised Spatial Analysis in Macroecology (SAM) Version 4.0 (Rangel et al., 2010) (available from https://www.ecoevol.ufg.br/sam/), R Version 4.0.5 (R Core Team, 2021), and Microsoft Excel. In order to run the semi-partial correlation test between adjusted X-chromosomal diversity (i.e., adjusted for distance from Africa) and minimum temperature, ppcor (Kim, 2015) was used initially. ppcor (Kim, 2015) was employed to assess whether adjusted mitochondrial diversity and minimum temperature are correlated.

As with Study 1, in parametric analysis, datapoints were highlighted as atypical when *z*-scored residuals surpassed |3.29| (e.g., Field, 2013). In instances where X-chromosomal diversity (SGDP data) was only available for one person (in a population), the diversity of that person was simply used to represent their population. In instances where the diversity of more than one person was available for a population, an average was calculated to represent their population. The same approach was used regarding autosomal diversity. As in Study 1, data were examined for the presence of positive spatial autocorrelation (Chen, 2016; Savin & White, 1977). However, positive spatial autocorrelation was not tested for when examining whether X-chromosomal diversity is correlated with distance from Africa. Instead, SAM Version 4.0, which adjusts the degrees of freedom (Rangel et al., 2010), was cautiously used in the event of spatial autocorrelation being present. Given the impact of spatial autocorrelation on Type I errors (Deblauwe et al., 2012), positive spatial autocorrelation was not tested for regarding whether there is an association between the ratio of X-chromosomal to autosomal diversity and climate, or when examining if mitochondrial diversity correlates with distance from Africa (given the results of the correlation tests). A Bonferroni correction (see Abdi, 2010) was applied.

## Results

### Study 1: Genetic and cranial diversities

#### Outside of Africa

For populations outside of Africa, X-chromosomal microsatellite heterozygosity (adjusted for distance from Africa) positively correlated with minimum temperature, s*r*(40) = .40, *p* = .040, spatial *DW* = 1.91. This result was found following Surui not being included due to their atypical standardised residual (*z* = -3.70) (as in part of Figure 2A). Adjusted autosomal SNP haplotype heterozygosity had no correlation with minimum temperature, s*r*(41) = .10, *p* = 1.00, spatial *DW* = 1.71. The same can be said for adjusted autosomal microsatellite heterozygosity, s*r*(39) = .25, *p* = .57, spatial *DW* = 1.92 (in the absence of Surui due to their standardised residual, *z* = -3.41). As for adjusted cranial form diversity, it was not found to have a correlation with minimum temperature for females, s*r*_*s*_(31) = -.13, *p* = 1.00, or for males, s*r*_*s*_(86) = -.02, *p* = 1.00.

#### Worldwide

X-chromosomal microsatellite heterozygosity (adjusted for distance from Africa) was found to have a positive association with minimum temperature, s*r*(46) = .39, *p* = .031, spatial *DW* = 1.92. However, there were caveats, with this correlation being calculated in the absence of two populations. When using 51 populations, Surui had an atypically low standardised residual (*z* = -3.52). Whether with or without Surui, heteroscedasticity was indicated. In the absence of Surui, out of the 50 remaining populations, any one of the populations was removed to see if any particular population may be driving the heteroscedasticity amongst the 50 populations. The absence of the Mbuti Pygmy population (Balloux et al., 2009) seemed to (visually) have the most impact. Indeed, without Mbuti Pygmy, heteroscedasticity appeared to be absent; amongst the 50 populations, it seemed like Mbuti Pygmy may have resulted in the heteroscedasticity. And so, without Surui and Mbuti Pygmy, there was a correlation between adjusted X-chromosomal diversity and minimum temperature (also see Supplementary Note 7). If a less stringent criterion (e.g., |1.96|) was used for detecting atypical datapoints (e.g., Field, 2013), then, when 51 populations were used, Mbuti Pygmy would have been atypical (*z* = -2.91).

Autosomal SNP haplotype heterozygosity (adjusted for distance from Africa) was not observed to be associated with minimum temperature, s*r*(48) = .16, *p* = 1.00, spatial *DW* = 1.81. Likewise, autosomal microsatellite heterozygosity (controlling for distance from Africa) did not yield a correlation with minimum temperature, s*r*(46) =.28, *p* =.28, spatial *DW* = 1.92 (without Surui because of their standardised residual, *z* = -3.69), and neither did Y-chromosomal microsatellite heterozygosity (adjusted for distance from Asia), s*r*_s_(48) = -.11, *p* = 1.00. A semi-partial Spearman test was used regarding Y-chromosomal diversity because heteroscedasticity seemed apparent in parametric analysis, and it persisted when any one population was absent (see Supplementary Note 8).

### Study 2: Diversity and diversity ratios

#### Diversity

When it came to the relationship between X-chromosomal diversity (SGDP data) and distance from Africa, two datapoints were indicated to be atypical. These datapoints were for the Mixtec population and the Palestinian population (Mallick et al., 2016); the former was atypically high (z = 3.62), and the latter was atypically low (z = -4.59). Therefore, those two datapoints were not used when i) testing whether X-chromosomal diversity is associated with distance from Africa, and when ii) X-chromosomal diversity was adjusted for distance from Africa. Congruent with previous research (Balloux et al., 2009), X-chromosomal heterozygosity (SGDP data) was found to decrease as distance from Africa advanced, *r*(12.30) = -.77, *p* = .006. Therefore, the X-chromosomal diversity (of populations who are located outside of Africa) was adjusted for distance from Africa. When examining if there is a correlation between adjusted X-chromosomal diversity and minimum temperate (i.e., if there is a significant semi-partial Pearson correlation), the author was unable to generate a *p*-value through the ppcor package. Therefore, the *p*-value was calculated from a *t*-value, and the *t*-value was arrived at (from the correlation coefficient) via a formula which is presented in Kim (2015). In contrast to results in Study 1, adjusted X-chromosomal diversity was not found to correlate with minimum temperature, s*r*(72) = -.04, *p* = 1.00 (Figure 3).

Previous research observed that mitochondrial diversity falls as populations are located at increasingly greater distances away from Africa (Balloux et al., 2009). In Study 2, distances between an origin (coordinates from von Cramon Taubadel & Lycett, 2008) and populations (Cenac, 2022) were used to check whether there is a correlation between distance from Africa and mitochondrial diversity. A correlation was not found, *r*(42) = -.37, *p* = .079. However, given that the decrease in genetic diversity is broadly found in genetic diversities, including mitochondrial previously (e.g., Balloux et al., 2009), it seems likely that the absence of a significant correlation in Study 2 was because analysis in Study 2 was underpowered. Therefore, analysis still (conservatively) occurred with mitochondrial diversity being adjusted for distance. When examining whether mitochondrial diversity (adjusted for distance from Africa) correlates with minimum temperature, heteroscedasticity was indicated. Therefore, a semi-partial Spearman’s test was used. Adjusted mitochondrial diversity was not found to correlate with minimum temperature, s*r*_s_(41) = .02, *p* = 1.00.

#### Diversity ratios

Research on diversity and climate has considered sex-biased migration (Balloux et al., 2009). It is unlikely that an association between mitochondrial diversity and climate is because of some relationship between climate and sex-biased migration—previous research did not find that the ratio of diversity between sex chromosomes is related to minimum temperature (Balloux et al., 2009). Therefore, it could seem improbable that the climatic signal in the X chromosome (Study 1) arose from an association between sex-biased migration and climate. However, Y-chromosomal diversity does not have a climatic signal (Balloux et al., 2009; Study 1), unlike X-chromosomal diversity in Study 1. Therefore, it would not be surprising if a link is actually indicated between sex-biased migration and climate. Although Balloux et al. (2009) examined if a link is indicated, they do not appear to have accounted for the expansion signal in X-chromosomal diversity; such an accounting could allow for a more powerful test of whether the sex chromosome diversity ratio is associated with climate. In Study 2, using HGDP-CEPH data (Balloux et al., 2009), the ratio of adjusted X-chromosomal to Y-chromosomal diversity increased with minimum temperature (Mbuti Pygmy and Surui were not included in analysis—see Study 1 results), *r*(47) = .45, *p* = .007, Spatial *DW* = 1.91; as minimum temperature cools, the X chromosome becomes less diverse relative to the Y chromosome.

When SGDP data (Mallick et al., 2016) were used, and the ratio of X-chromosomal to autosomal diversity was examined (with neither diversity being adjusted for distance from Africa), analysis did not find a significant correlation between the ratio and minimum temperature, *r*(71) = - .04, *p* = 1.00. Pima and Surui populations (Mallick et al., 2016) were not used in the correlation analysis due to their atypical residuals (*z*s = -3.40 and -4.57 respectively). Mixtec and Palestinian populations (Mallick et al., 2016) were not used due to the atypicality (mentioned above) of their diversities for their distance from Africa.

## Discussion

Previous research has not found support for a climatic signal being present in autosomal diversity, Y-chromosomal diversity (Balloux et al., 2009), or cranial form diversity (Betti et al., 2009). This lack of support was matched in the present manuscript which addressed potential limitations of previous studies (see *Introduction*). On the other hand, Study 1 diverged from previous research (i.e., Balloux et al., 2009) when it came to X-chromosomal diversity; in Balloux et al. (2009), a climatic signal was not indicated in X-chromosomal diversity, unlike in Study 1.

This incongruity between Balloux et al. (2009) and Study 1 is unlikely to have been caused by Balloux et al. and Study 1 not using the same location as the *origin*—similar origins were used regarding X-chromosomal diversity (*Introduction*; Table S1). Instead, the incongruity likely arose because correlation tests in Study 1 omitted populations (Mbuti Pygmy and Surui) which Balloux et al. (2009) included in their analysis. In the present research, if the correlation coefficient is calculated when those two populations are included (global analysis), the coefficient is numerically smaller in magnitude (*r* = .18) than when the populations are absent. If the coefficient is calculated without Surui, but including Mbuti Pygmy, the correlation coefficient is also numerically smaller (*r* = .28). It is also worth stating that when only populations outside of Africa are used, the inclusion of Surui (who had an atypical standardised residual) leads to a numerically weaker correlation coefficient (*r* = .31) than when Surui are absent (Figure 2A).

In contrast to Study 1, Study 2 did not lend support to there being a climatic signal in X-chromosomal diversity. Therefore, the present research leaves a question mark when answering if there is a climatic signal. Regarding X-chromosomal diversity, more populations were used in Study 2 (SGDP data) than in Study 1 (HGDP-CEPH data) (Balloux et al., 2009; Mallick et al., 2016; see *Method*). However, per population, far fewer individuals were represented in the SGDP data (see Balloux et al., 2009; Mallick et al., 2016; Table S2); heterozygosity from the HGDP-CEPH data may generally be more representative of populations. Therefore, it could be reasonable to *provisionally* favour results found with the HGDP-CEPH data (i.e., Study 1). However, further research clearly is required in order to say, with any reasonable confidence, if there is a climatic signal.

Mitochondria are recognised for being tied to climatic adaptation (Grover-Thomas et al., 2026), unlike the X chromosome (e.g., Lasne et al., 2019). Therefore, Study 1’s indication of a climatic signal in the X chromosome may be surprising. Nevertheless, there are, theoretically, other routes through which genetic diversity could be related to climate, distinct of climatic adaptation, e.g., a link between sex-biased migration and climate (Balloux et al., 2009). However, previous research did not find a relationship between the sex chromosome diversity ratio and climate, and this absence of a relationship suggested that the climatic signal in mitochondrial diversity is not explainable by a relationship between sex-biased migration and climate (Balloux et al., 2009). On the other hand, Study 2 (using HGDP-CEPH data) found a relationship between the ratio and climate; that finding may give weight to the idea that the climatic signal in mitochondrial diversity *and* X-chromosomal diversity could reflect an association between climate and sex-biased migration. Of course, given the apparent climatic signal in X-chromosomal diversity when HGDP-CEPH data are used, perhaps it is reasonable to expect a relationship between the sex chromosome diversity ratio (calculated from HGDP-CEPH data) and climate in Study 2 (see *Method*). Consequently, because a climatic signal in the X chromosome was not evident in SGDP data (Study 2), there is clear reason to question the correlation between the ratio and climate. Moreover, when SGDP data were used, an association was not observed between the ratio of X-chromosomal to autosomal diversity and climate. Could it be that this absence of an association was because X-chromosomal (or autosomal) diversity was not adjusted for distance from Africa? The answer likely is *no*—that is because there does not seem to be a climatic signal in X-chromosomal diversity using SGDP data (Figure 3).

Given the association between mitochondrial diversity and climate (with two different datasets regarding mitochondrial diversity) (Balloux et al., 2009), it is surprising that the present research did not indicate a climatic signal in mitochondrial diversity. Given the connection between mitochondria and climatic adaptation (Grover-Thomas et al., 2026), and Balloux et al. (2009), it seems likely that the absence of a climatic signal in mitochondrial diversity in Study 2 is an exception to the norm.

When examining if adjusted diversity is related to minimum temperature, heteroscedasticity i) may have been present in some analyses in Balloux et al. (2009) (see Supplementary Note 8), and it ii) seems to have occurred in the present research when analysis used adjusted X-chromosomal diversity (50 populations) and adjusted Y-chromosomal diversity in Study 1, and when analysis used mitochondrial diversity in Study 2. Heteroscedasticity is known to arise for a number of reasons, e.g., crucial variables not being included in analysis (e.g., Ohaegbulem & Iheaka, 2024); research on climate and diversity has not been limited to minimum temperature when it comes to climatic variables (Betti et al., 2009, 2012; Manica et al., 2007). Therefore, the actual/potential instances of heteroscedasticity in Balloux et al. and Studies 1 and 2 could possibly have arisen because certain climatic variables were not taken into account. Consequently, regarding climatic variables, it would seem to be prudent for future research to look beyond minimum temperature when addressing if there is a climatic signal in X-chromosomal and mitochondrial diversities. Nevertheless, in previous research which found a relationship between climate and the diversities of the tibia and femur, backward stepwise regression resulted in minimum temperature being included as the only predictor/independent variable (not maximum temperature, precipitation, climatic interaction terms, or indeed distance from Africa) regarding the prediction of diversity (Betti et al., 2012).

## Conclusion

In agreement with previous research (Balloux et al., 2009; Betti et al., 2009), a climatic signal did not seem to be suggested in autosomal, Y-chromosomal, or cranial form diversities. In contrast to previous research (Balloux et al., 2009), a climatic signal i) was not indicated in mitochondrial diversity, and ii) was indicated in X-chromosomal diversity in Study 1. However, no such signal was indicated in Study 2 with respect to X-chromosomal diversity. On balance (Balloux et al., 2009; Grover-Thomas et al., 2026; Study 2), it seems probable that there *is* a climatic signal in mitochondrial diversity (in general). Given the conflicting findings over whether there is a climactic signal in X-chromosomal diversity (Studies 1 and 2), it seems rather questionable if there truly is a climatic signal in the X chromosome. Hopefully, the present research i) functions as a reminder of how important replicability is (see Nosek, & Errington, 2020) and, ii) encourages further replication research on whether X-chromosomal diversity has a climatic signal.

