## Supplementary material for "X-chromosomal diversity may, or may not, reflect climate"

**Supplementary material: X-chromosomal diversity may, or may not, reflect climate**

**Supplementary notes**

**Supplementary Note 1**. Diversity was adjusted for distance by fitting a linear trend (from the relationship between diversity and distance), and calculating residuals (i.e., the residuals were diversity adjusted for linear distance) (Balloux et al., 2009).

**Supplementary Note 2**. The autosomal diversities were microsatellite heterozygosity and SNP haplotype heterozygosity, and, for the X and Y chromosomes, diversities were microsatellite heterozygosity (Balloux et al., 2009).

**Supplementary Note 3**. Nonetheless, for a number of cranial dimensions, their diversity is associated with climate (Manica et al., 2007).

**Supplementary Note 4**. Several diversities were of interest in Study 1. These were: autosomal heterozygosity (microsatellite, and SNP haplotype separately), X-chromosomal microsatellite heterozygosity, Y-chromosomal microsatellite heterozygosity, and cranial form diversity. Each undergoes a decline with the greatening of distance from Africa (Balloux et al., 2009; Betti et al., 2009; Cenac, 2022; Prugnolle et al., 2005). Genetic diversities which are consistent with global expansion originating from Africa (alone) are autosomal microsatellite heterozygosity (Manica et al., 2007), autosomal SNP haplotype heterozygosity, and X-chromosomal microsatellite heterozygosity, whereas, Y-chromosomal microsatellite heterozygosity seemingly corresponds to expansion from elsewhere (Cenac, 2022). As for cranial form diversity, when the diversity is of male crania, an expansion is indicated to have begun in Africa alone (Betti et al., 2009; Cenac, 2022). With respect to the cranial form diversity of females, Betti et al. (2009) found the area from which the expansion likely originated was an area that was not utterly in Africa. However, in a preprint, the area was in Africa in its entirety (Cenac, 2022). Interestingly, cranial form diversity appears to lead to a larger area of potential origin than cranial shape diversity does (Cenac, 2022). Therefore, the area in Betti et al. (2009) likely would have been in Africa alone if they had used cranial shape diversity (given that the peak point was in sub-Saharan Africa in their study). Hence, whilst the area of origin was not exclusive to Africa in Betti et al. for the form diversity of female crania, it still makes sense to adjust the cranial form diversity of females for distance from Africa in order to account for the expansion signal.

**Supplementary Note 5**. In addition to minimum temperature, Betti et al. (2009, 2012) and Manica et al. (2007) also included other climatic variables in their analyses.

**Supplementary Note 6**. However, without two populations (both Patagonian), there was a linear relationship (Betti et al., 2009)—see Betti et al. (2009) for discussion about this.

**Supplementary Note 7**. The intention was to adjust diversity for distance from the peak point. The peak point used with respect to X-chromosomal diversity was found by using 51 populations (Cenac,

2022). Peak points may be substantially influenced by African populations (Cenac, 2022). So, without Mbuti Pygmy (and Surui, i.e., using 49 populations), X-chromosomal diversity may generate a markedly different peak point than when 51 populations are used. The location of the peak point does likely affect the correlation between diversity (adjusted for distance) and minimum temperature (Figure 2). When using 49 populations, the peak point was more southern, with its location being (30°S, 20°E).[1] When X-chromosomal diversity was adjusted for distance from (30°S, 20°E) and 51 populations were used, Surui still had a very low standardised residual ($z$ = -3.56), and, in the absence of Surui, Mbuti Pygmy seemed to still be driving heteroscedasticity. Using 49 populations (i.e., not using Surui or Mbuti Pygmy), the correlation coefficient for the relationship between X-chromosomal diversity [adjusted for distance from (30°S, 20°E)] and minimum temperature was, $r$ = .39, i.e., the same (to two decimal places) as what was found when using the peak point calculated from the 51 populations. And so, no matter whether the peak point for 51 or 49 populations was used, the correlation coefficient did not seem to be notably affected.

**Supplementary Note 8**. Figure 3 in Balloux et al. (2009) may indicate heteroscedasticity with respect to parametrically testing for relationships between heterozygosity (adjusted for distance from Africa) and minimum temperature when the heterozygosity used was of X-chromosomal microsatellites, autosomal microsatellites, autosomal SNP haplotypes, and Y-chromosomal microsatellites (in addition to regression analyses, Balloux et al. employed partial Mantel tests).

---

[1] Distances calculated in a preprint (Cenac, 2022) were used. These were distances between coordinates in the continent of Africa from Betti et al. (2013) and HGDP-CEPH populations (Cenac, 2022). In a preprint, those distances were used to calculate peak points (Cenac, 2022), hence why the distances were used in the current research.

### Supplementary tables

**Table S1**

*Peak Points*

| Diversity | Peak point | Source of peak point |
|---|---|---|
| Autosomal STR heterozygosity | 25°S, 15°E | Cenac (2023) |
| Autosomal SNP haplotype heterozygosity | 15°N, 5°E | Cenac (2022) |
| X-chromosomal STR heterozygosity | 15°S, 15°E | Cenac (2022) |
| Y-chromosomal STR heterozygosity | Delhi, India | von Cramon and Lycett (2008)—see that article for coordinates |
| Mitochondrial diversity | 25°S, 15°E | Cenac (2022) |

*Note*. See Table S3 for distances from the peak points (Cenac, 2022).

**Table S2**

*Statistics for Populations*

| Population | Sample size | | Minimum temperature (°C) | Adjusted X-chromosomal heterozygosity | Ratio: X-chromosomal diversity divided by autosomal diversity | Distance from Africa (km) |
|---|---|---|---|---|---|---|
| | X chromosome | Autosome | | | | |
| Adygei | 1 | 2 | -3.51 | -0.00001869 | 0.5753 | 7279.77 |
| Albanian | 1 | 1 | -10.80 | -0.00002510 | 0.5674 | 7567.27 |
| Aleut | 1 | 2 | -8.82 | 0.00007425 | 0.6172 | 15180.38 |
| Australian | 1 | 2 | 14.77 | 0.00003959 | 0.6015 | 18335.90 |
| Basque | 1 | 2 | -10.27 | -0.00001051 | 0.5775 | 9192.67 |
| BedouinB | 1 | 2 | 0.79 | -0.00002295 | 0.5931 | 5914.21 |
| Bengali | 1 | 2 | 8.67 | 0.00004011 | 0.5875 | 11377.81 |
| Bergamo | 1 | 2 | -17.64 | -0.00000136 | 0.5702 | 8429.96 |
| Bougainville | 2 | 2 | 23.05 | -0.00000245 | 0.5467 | 19223.37 |
| Burusho | 1 | 2 | -26.16 | 0.00005064 | 0.6207 | 9543.94 |
| Cambodian | 1 | 2 | 16.86 | 0.00002054 | 0.5848 | 13319.46 |
| Chipewyan | 1 | 2 | -47.27 | -0.00006603 | 0.5132 | 18489.92 |
| Cree | 1 | 2 | -37.79 | 0.00004051 | 0.5502 | 19441.13 |
| Dai | 2 | 4 | 4.49 | -0.00000317 | 0.5852 | 12404.60 |
| Daur | 1 | 1 | -33.15 | 0.00002537 | 0.5949 | 13300.22 |
| Druze | 1 | 2 | 2.13 | 0.00002254 | 0.6363 | 5968.21 |
| Dusun | 2 | 2 | 20.05 | -0.00001755 | 0.5655 | 14644.94 |
| English | 1 | 2 | -2.25 | 0.00002001 | 0.6099 | 9250.42 |
| Eskimo Naukan | 2 | 2 | -44.01 | -0.00002272 | 0.5813 | 14347.95 |
| Eskimo Sireniki | 1 | 2 | -44.87 | 0.00000010 | 0.5868 | 14606.89 |
| Even | 2 | 3 | -48.93 | 0.00000555 | 0.5759 | 13598.27 |
| Finnish | 1 | 3 | -15.89 | -0.00001502 | 0.5756 | 9024.70 |
| French | 1 | 3 | -9.13 | -0.00000590 | 0.5848 | 9037.99 |
| Han | 1 | 3 | -9.24 | 0.00001349 | 0.5825 | 13201.97 |
| Hezhen | 1 | 2 | -37.46 | 0.00000652 | 0.5771 | 13985.20 |
| Hungarian | 1 | 2 | -14.48 | -0.00003220 | 0.5510 | 7891.02 |
| Icelandic | 2 | 2 | -11.46 | 0.00000128 | 0.5769 | 10950.94 |
| Igorot | 1 | 2 | 12.55 | -0.00002132 | 0.5752 | 15227.86 |
| Iraqi Jew | 1 | 2 | -2.68 | -0.00001669 | 0.5844 | 6834.25 |
| Itelman | 1 | 1 | -25.60 | 0.00001855 | 0.6320 | 14656.18 |
| Japanese | 1 | 3 | -1.06 | -0.00000778 | 0.5388 | 14842.96 |
| Kalash | 1 | 2 | -30.77 | 0.00000765 | 0.6153 | 9321.50 |
| Karitiana | 1 | 3 | 14.96 | 0.00006162 | 0.6348 | 27360.70 |
| Kinh | 1 | 2 | 3.64 | -0.00001965 | 0.5520 | 12963.45 |
| Korean | 1 | 2 | -15.19 | 0.00002692 | 0.5950 | 14029.38 |
| Kyrgyz | 1 | 2 | -12.48 | -0.00003663 | 0.5342 | 9624.69 |
| Lahu | 1 | 2 | 2.76 | -0.00005307 | 0.5271 | 12360.60 |
| Makrani | 1 | 2 | 0.71 | -0.00001733 | 0.5745 | 8766.88 |
| Mala | 1 | 3 | 18.46 | 0.00006928 | 0.6258 | 10950.89 |

| | | | | | | |
|---|---|---|---|---|---|---|
| Mansi | 1 | 2 | -41.84 | 0.00003980 | 0.6279 | 9894.89 |
| Mayan | 2 | 2 | 17.73 | 0.00002233 | 0.5704 | 22929.18 |
| Miao | 1 | 2 | -6.41 | 0.00001214 | 0.5965 | 12940.11 |
| Mixe | 3 | 3 | 0.71 | 0.00001823 | 0.5641 | 22862.38 |
| Mongola | 1 | 2 | -27.78 | 0.00001309 | 0.5858 | 12489.00 |
| Naxi | 1 | 3 | -4.18 | -0.00002556 | 0.5287 | 12193.97 |
| Onge | 2 | 2 | 24.67 | -0.00007899 | 0.5544 | 12146.23 |
| Orcadian | 1 | 2 | 0.88 | 0.00001209 | 0.5912 | 9816.67 |
| Oroqen | 1 | 2 | -39.44 | -0.00004987 | 0.5320 | 13385.16 |
| Papuan | 3 | 15 | 21.35 | -0.00002200 | 0.5790 | 17898.87 |
| Pathan | 1 | 2 | -3.46 | 0.00005695 | 0.6280 | 9244.22 |
| Piapoco | 2 | 2 | 20.24 | 0.00001507 | 0.5548 | 26018.88 |
| Pima | 1 | 2 | -6.51 | -0.00009833 | 0.4359 | 21129.89 |
| Punjabi | 2 | 4 | 1.85 | 0.00000786 | 0.5842 | 9624.06 |
| Quechua | 2 | 3 | -2.30 | 0.00006017 | 0.5537 | 27087.94 |
| Russian | 1 | 2 | -35.80 | 0.00002967 | 0.6123 | 9101.15 |
| Saami | 1 | 2 | -27.69 | -0.00005043 | 0.5418 | 10092.49 |
| Sardinian | 1 | 3 | -1.61 | 0.00005415 | 0.6611 | 8486.69 |
| She | 1 | 2 | -7.25 | -0.00000339 | 0.5716 | 13883.76 |
| Sherpa | 1 | 2 | -24.51 | -0.00004134 | 0.5735 | 10882.01 |
| Sindhi | 1 | 2 | 4.69 | 0.00001612 | 0.5978 | 9263.29 |
| Spanish | 1 | 2 | -5.28 | 0.00004191 | 0.6224 | 9574.09 |
| Surui | 2 | 2 | 13.85 | -0.00011648 | 0.3834 | 27516.53 |
| Thai | 1 | 2 | 13.62 | -0.00003317 | 0.5285 | 12794.68 |
| Tibetan | 2 | 2 | -28.70 | -0.00002267 | 0.5733 | 10623.03 |
| Tlingit | 1 | 2 | -20.20 | 0.00005304 | 0.5813 | 15044.72 |
| Tu | 1 | 2 | -20.40 | 0.00000114 | 0.5739 | 11937.53 |
| Tubalar | 2 | 2 | -38.19 | 0.00003168 | 0.6181 | 10661.32 |
| Tujia | 1 | 2 | -6.06 | -0.00001424 | 0.5740 | 12897.90 |
| Turkish | 1 | 2 | -16.70 | -0.00001757 | 0.5830 | 6624.68 |
| Tuscan | 1 | 2 | -4.32 | 0.00000247 | 0.6029 | 8299.38 |
| Ulchi | 2 | 2 | -38.20 | -0.00003115 | 0.5406 | 14133.05 |
| Uygur | 1 | 2 | -17.19 | 0.00003708 | 0.5960 | 10149.32 |
| Yakut | 1 | 2 | -55.28 | 0.00002892 | 0.6124 | 13027.38 |
| Yemenite Jew | 1 | 2 | 1.17 | -0.00004835 | 0.5269 | 7651.32 |
| Yi | 1 | 2 | -8.79 | -0.00002878 | 0.5564 | 12392.77 |

*Note*. Population names and sample sizes were obtained from Mallick et al. (2016). Data from NASA POWER (2026) were used to calculate the minimum temperatures (which are averages). X-chromosomal diversity (Mallick et al., 2016) was adjusted for distance from Africa. X-chromosomal diversities and autosomal diversities from previous research (Mallick et al., 2016) were used to calculate the ratios. The distances presented in Table S2 are the ones which were used to adjust X-chromosomal diversity. Distances were from coordinates given in von Cramon-Taubadel & Lycett (2008). Those coordinates are for a location in southern Africa (von Cramon-Taubadel & Lycett, 2008). Von Cramon-Taubadel and Lycett (2008) calculated distances from that location, as well as from other locations. Southern Africa was selected for use in Study 2 because, amongst regions of

Africa, the south seems to be the most probable region for being where the global expansion began (Cenac, 2022). Nonetheless, because the analysis in Study 2 (on whether X-chromosomal diversity has a climatic signal) was restricted to populations that are located outside of Africa (Mallick et al., 2016), it does not matter which (continental) African location was employed as the origin (Figure 2A).

**Table S3**

*Distances from Locations for HGDP-CEPH Populations*

| Population | Distance (km) from (25°S, 15°E) | Distance (km) from (15°N, 5°E) | Distance (km) from (15°S, 15°E) | Distance (km) from Delhi, India |
|---|---|---|---|---|
| Adygei | 8047.95 | 4894.27 | 6995.06 | 3771.47 |
| Balochi | 9673.48 | 6519.80 | 8620.59 | 1039.09 |
| Bantu Kenya | 3393.53 | 4049.17 | 2753.78 | 8043.01 |
| Basque | 9960.85 | 6807.17 | 8907.97 | 6885.60 |
| Bedouin | 6682.40 | 3528.72 | 5629.51 | 4037.60 |
| Biaka Pygmy | 3229.71 | 1793.81 | 2122.73 | 7628.69 |
| Brahui | 9673.48 | 6519.80 | 8620.59 | 1039.09 |
| Burusho | 10312.13 | 7158.45 | 9259.24 | 932.21 |
| Cambodian | 14087.64 | 10933.96 | 13034.75 | 3431.93 |
| Colombian | 26787.07 | 23633.39 | 25734.18 | 18938.61 |
| Dai | 13172.79 | 10019.11 | 12119.90 | 2460.91 |
| Daur | 14068.41 | 10914.73 | 13015.52 | 4557.82 |
| Druze | 6736.40 | 3582.72 | 5683.51 | 4025.28 |
| French | 9806.18 | 6652.50 | 8753.29 | 6658.06 |
| Han | 13755.22 | 10601.54 | 12702.33 | 3564.88 |
| Hazara | 9966.21 | 6812.53 | 8913.33 | 867.47 |
| Hezhen | 14753.39 | 11599.71 | 13700.50 | 5244.40 |
| Italian | 9198.15 | 6044.47 | 8145.26 | 6047.68 |
| Japanese | 15611.14 | 12457.47 | 14558.26 | 5665.59 |
| Kalash | 10089.69 | 6936.01 | 9036.80 | 980.21 |
| Karitiana | 28128.88 | 24975.20 | 27076.00 | 20280.43 |
| Lahu | 13128.78 | 9975.10 | 12075.90 | 2417.07 |
| Makrani | 9535.07 | 6381.39 | 8482.18 | 1313.20 |
| Mandenka | 5046.10 | 1866.24 | 4222.39 | 9282.07 |
| Maya | 23697.37 | 20543.69 | 22644.48 | 15848.91 |
| Mbuti Pygmy | 3258.32 | 3058.71 | 2351.34 | 7575.79 |
| Melanesian | 19991.55 | 16837.87 | 18938.66 | 9361.43 |
| Miao | 13708.29 | 10554.61 | 12655.40 | 3123.44 |
| Mongola | 13723.86 | 10570.18 | 12670.97 | 4200.70 |
| Mozabite | 6460.53 | 1899.88 | 5376.75 | 7100.66 |
| Naxi | 12962.16 | 9808.48 | 11909.27 | 2285.50 |
| Orcadian | 10584.86 | 7431.18 | 9531.97 | 6757.97 |
| Oroqen | 14148.96 | 10995.28 | 13096.07 | 4779.35 |
| Palestinian | 6736.40 | 3582.72 | 5683.51 | 4025.28 |
| Papuan | 18667.05 | 15513.37 | 17614.17 | 8036.93 |
| Pathan | 10012.40 | 6858.73 | 8959.52 | 831.49 |
| Pima | 21898.07 | 18744.39 | 20845.18 | 14049.61 |
| Russian | 9869.34 | 6715.66 | 8816.45 | 4527.82 |
| San | 677.45 | 4322.09 | 850.52 | 10148.32 |
| Sardinian | 9254.88 | 6101.20 | 8201.99 | 6224.96 |

| | | | | |
|---|---|---|---|---|
| She | 14651.94 | 11498.26 | 13599.05 | 4112.54 |
| Sindhi | 10031.47 | 6877.79 | 8978.58 | 859.09 |
| Surui | 28284.72 | 25131.04 | 27231.83 | 20436.26 |
| Tu | 12705.72 | 9552.04 | 11652.83 | 2396.71 |
| Tujia | 13666.09 | 10512.41 | 12613.20 | 3108.38 |
| Tuscan | 9067.57 | 5913.89 | 8014.68 | 6007.90 |
| Uygur | 10917.51 | 7763.83 | 9864.62 | 1758.67 |
| Xibo | 10955.08 | 7801.40 | 9902.19 | 1714.47 |
| Yakut | 13795.57 | 10641.89 | 12742.68 | 5343.17 |
| Yi | 13160.95 | 10007.27 | 12108.06 | 2540.62 |
| Yoruba | 3823.40 | 777.84 | 2783.04 | 8062.29 |

*Note*. The names of populations were obtained from Balloux et al. (2009), and distances are from Cenac (2022).
